# Reverse-transcription recombinase-aided amplification assay for H5 subtype avian influenza virus

**DOI:** 10.1101/2020.10.22.350025

**Authors:** Suchun Wang, Yang Li, Nan Jiang, Fuyou Zhang, Qingye Zhuang, Guangyu Hou, Lijian Jiang, Jianmin Yu, Xiaohui Yu, Hualei Liu, Chenglong Zhao, Liping Yuan, Baoxu Huang, Kaicheng Wang

## Abstract

The H5 subtype Avian Influenza Virus has caused huge economic losses to the poultry industry and is a threat to human health. A rapid and simple test is needed to confirm infection in suspected cases during disease outbreaks. In this study, we developed a reverse-transcription recombinase-aided amplification assay for the detection of H5 subtype avian influenza virus. Assays were performed at a single temperature (39°C), and the results were obtained within 20 min. The assay showed no cross-detection with Newcastle disease virus or infectious bronchitis virus. The analytical sensitivity was 10^3^ RNA copies per reaction at a 95% confidence interval according to probit regression analysis, with 100% specificity. Compared with published reverse-transcription quantitative real-time polymerase chain reaction assays, the κ value of the reverse transcription recombinase-aided amplification assay in 365 avian clinical samples was 0.970 (p < 0.001). The sensitivity for avian clinical sample detection was 94.44% (95%CI, 70.63% - 99.71%), and the specificity was 100% (95%CI, 98.64% - 100%). These results indicated that our reverse-transcription recombinase-aided amplification assay may be a valuable tool for detecting H5 subtype avian influenza virus.

## INTRODUCTION

Avian influenza virus (AIV) is a negative-sense RNA virus that belongs to the orthomyxoviridae family [1]. The AIV genome is composed of eight distinct RNA segments encoding at least 10 proteins which coordinate functions, components and structure of the virus[2,3]. AIV can be classified into highly pathogenic avian influenza virus (HPAIV) and low pathogenic avian influenza virus (LPAIV) based on pathogenicity in chickens. Since the first H5N1 HPAIV was detected in 1996, these viruses have been prevalent among poultry in Asia, Europe, and Africa. The viruses constantly undergo genetic drift and shift that permanently threats poultry industry and human health[4].

All LPAIV and HPAIV infections of subtypes H5 in poultry are notifiable to the World Organization for Animal Health (O.I.E.). Determination of the type of AIV is of utmost importance for the diagnosis of these infections. This can be achieved biologically by determination of the intravenous pathogenicity index (IVPI) in experimentally inoculated chickens or molecularly by nucleotide sequence analysis of the site encoding the AIV. Since animal experiment facilities or expensive equipment are required for either pathway, solutions for alternative techniques have been sought in the past. These included restriction enzyme cleavage patterns, probe hybridization and real time RT-PCR (RT-qPCR) approaches [5–13]. Based on the widespread availability of RT-qPCR technology in diagnostic laboratories and its recent favorable use in pathotyping of H5 subtype HPAIV of the goose/Guangdong (gs/GD) lineage.

Recently, rapid isothermal amplification techniques have been developed, such as loop-mediated isothermal amplification (LAMP) [14], recombinase polymerase amplification (RPA) [15], recombinase assisted amplification (RAA) (16)and strand displacement amplification (SDA) [16], and used for large-scale testing. Among these rapid nucleic acid detection methods, reverse transcription recombinase-mediated isothermal amplification (RT-RAA) is a rapid thermostatic nucleic acid amplification technology that utilizes a recombinant enzyme obtained from bacteria or fungi. At normal temperature, the recombinant enzyme can Tightly bind to the primer DNA to form a polymer of enzymes and primers. When the primer searches the template DNA for a complementary sequence that perfectly matches it, with the help of a single-stranded DNA binding protein, open the double-stranded structure of the template DNA Under the action of DNA polymerase, a new complementary DNA strand is formed, and the amplification product grows exponentially. This technology has the characteristics of high sensitivity, stronger specificity and reliability.

In this study, in order to efficiently detect H5 subtype AIV, a RT-RAA assay was designed and its analytical specificity and sensitivity were used to evaluate. Our study suggest that RT-RAA meets the need of field test, presents a rapid and sensitive detection method that can be used as an alternative to animal inoculation or nucleotide sequencing.

## Materials and methods

### Ethics statement

This study was conducted according to the animal welfare guidelines of the World Organization for Animal Health [17] and was approved by the Animal Welfare Committee of the China Animal Health and Epidemiology Center (CAHEC). CAHEC has permission to engage in activities of highly pathogenic avian influenza virus. The swab samples were collected for this study after being granted permission by multiple relevant parties, including the Ministry of Agriculture and Rural Affairs of China, the China Animal Health and Epidemiology Center, the relevant veterinary sections of the provincial and county governments, and the relevant farm owners.

### Samples and extraction of viral nucleic acids

All the AIV used in this study were isolated and identified in the National Avian Influenza Professional Laboratory in China Animal Health and Epidemiology Center and were stored at −80°C. The Newcastle disease virus (NDV) and infectious bronchitis virus (IBV) were all maintained at our lab. The samples were centrifuged at 12,000× g for 10 min and the supernatant from each sample was used for RNA extraction on the QIAxtractor platform using a cador Pathogen 96 QIAcube HT kit (Qiagen, Hilden) according to the manufacturer’s instructions. The extracted RNA were stored at −80°C for subsequent tests.

### Preparation of plasmid standard

A H5 subtype AIV plasmid standard was developed using thereference strain A/duck/Yunnan/5310/2006(H5N1) (GenBank accession number CY030889). A 1776 bp fragment of the whole hemagglutinin (HA) gene of H5 subtype AIV was cloned into the pUC57 vector to quantify DNA copy number. The recombinant plasmid DNA were prepared with a SanPrep Column Plasmid Mini-Preps Kit (Sangon) and quantified using a Thermo Scientific Multiskan GO Microplate Photometer (Thermo Fisher Scientific). The DNA copy number was calculated according to the following formula: DNA copy number = (copy number/μl) = [6.02 × 10^23^ ×plasmid concentration (ng/μl)×10^−9^]/[DNA length × 660]. The sensitivity of the RT-RAA was evaluated by real-time fluorescence detection, using a dilution series of standard recombinant plasmids ranging from 10^6^-10^1^ DNA copies per reaction.

### Design of H5 RT-RAA primers and exo-probes

To detect H5 subtype AIV, a total of 4636 available HA gene segments of H5 subtype AIV obtained from GenBank database were aligned, which contained the HA gene sequence of all currently circulating branches of H5 subtype AIV, and highly conserved regions were subsequently identified with Molecular Evolutionary Genetics Analysis (MEGA) software 6.0 [18, 19] for the design the gene-specific primers and probes. Primers were designed using OLIGO 7 software [20] and showed no major non-specific sequence similarities by BLAST analysis. Three H5 forward primers and five reverse primers were designed to select the best primers and probes in combination. The appropriate primers and probe used in this study were shown in Table 1. The 30th base at the 5’ end of the probe was labeled with the FAM light-emitting group. The 30th base was connected to the abasic site Tetrahydrofuran (THF). The 31st base was labeled with the BHQ1 quenching group, and the 3’ end was modified by C3-spacer blocking. All the primers and probes were synthesized by Sangon Biotech.

**TABLE 1.**
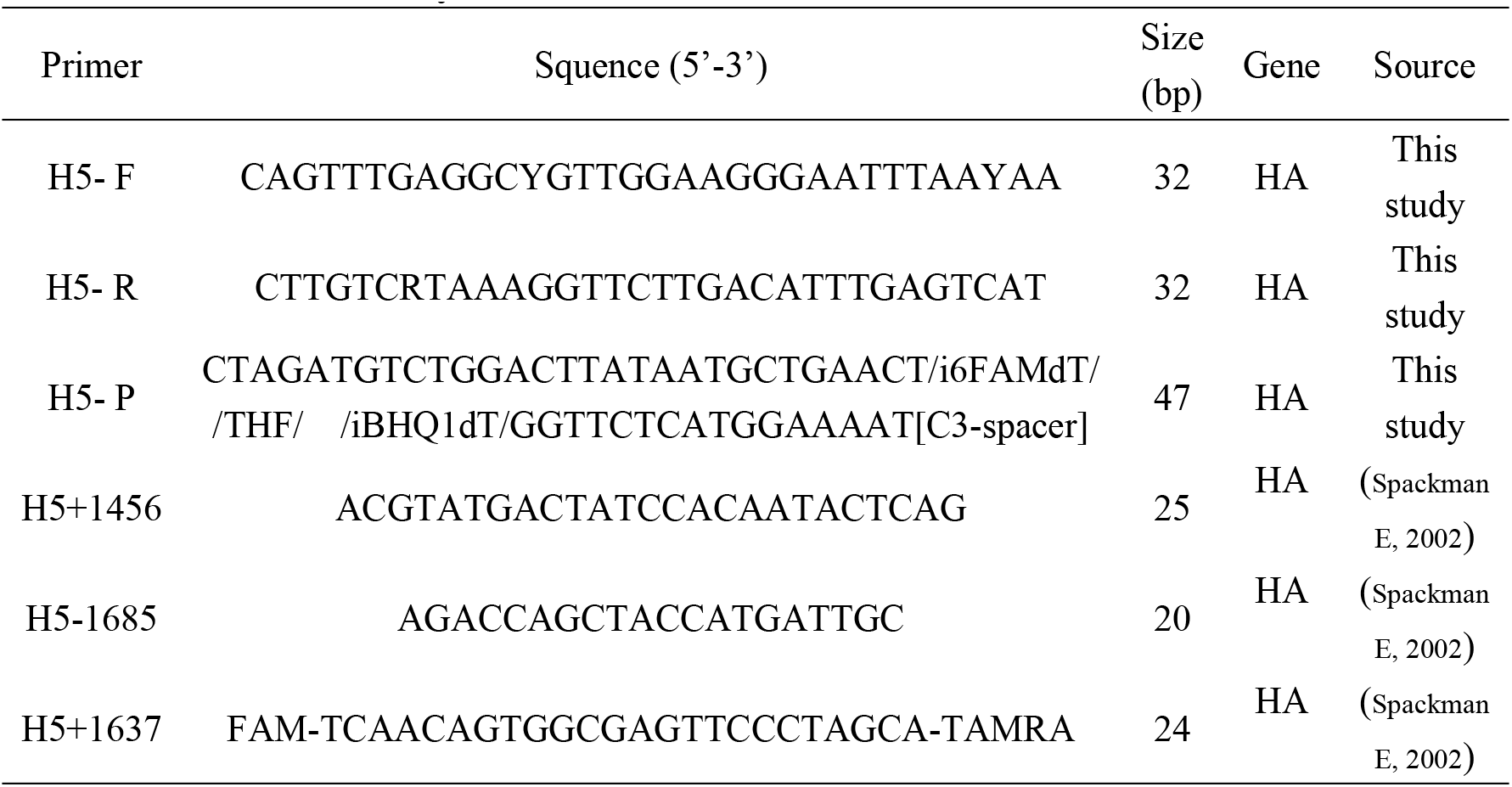
Primer and probe sequences used for RT-RAA, RT-qPCR and conventional PCR assays.

### RT-RAA for detection of H5 subtype AIV

The appropriate primers and exo-probes were screened by H5 RAA assay and verified by H5 RT-RAA assay. According to the manufacturer’s instructions, the RT-RAA reaction was performed with an RT exo kit in 50 μl reaction mixture including all the necessary enzymes and reagents for RT and DNA amplification in lyophilized pellets (Jiangsu Qitian Bio-Tech Co. Ltd.). The reaction mixture contained the following: 2 μl RNA template, 25 μl rehydration buffer, 15.7 μl ddH_2_O, 2.5 μl of magnesium acetate, 2.1 μl of each primer (10 μM) and 0.6 μl target-specific RT-RAA exo-probe. For amplification, the tubes were then transferred to a tube holder in an RT-RAA fluorescence detection device (QT-RAA-F7200; Jiangsu Qitian Bio-Tech Co. Ltd.) set at 39°C for 20 min. Each run included nuclease-free water as a negative control.

### Specificity, sensitivity and reproducibility of RT-RAA

Using the RNA of H5 subtype AIV as the template, a total of 7 groups of RNA with different concentrations were established for nucleic acid amplification under the optimal RT-RAA conditions. Then the selected appropriate primers and external probes were verified by H5 RT-RAA analysis. And the sensitivity was determined using serially diluted mixture of HA plasmids (each plasmid was 10^7^ copies/μL −10^1^ copies/μL) as quantitative standards. Then take 2 μL as the reaction template, and perform RT-RAA amplification according to the aforementioned loading method, with eight replicates for each dilution.

The specificity of the RT-RAA assay for H5 subtype AIV was evaluated using four H5-positive AIVs, 10 other subtype AIVs (H1N2, H3N2, H4N2, H6N2, H7N3, H7N9, H9N2, H10N7, H11N9), two NDVs and two IBVs. These viruses are the main respiratory viruses affecting birds and were previously identified by our lab. The details of all the viruses tested are listed in Table 2.

**TABLE 2.**
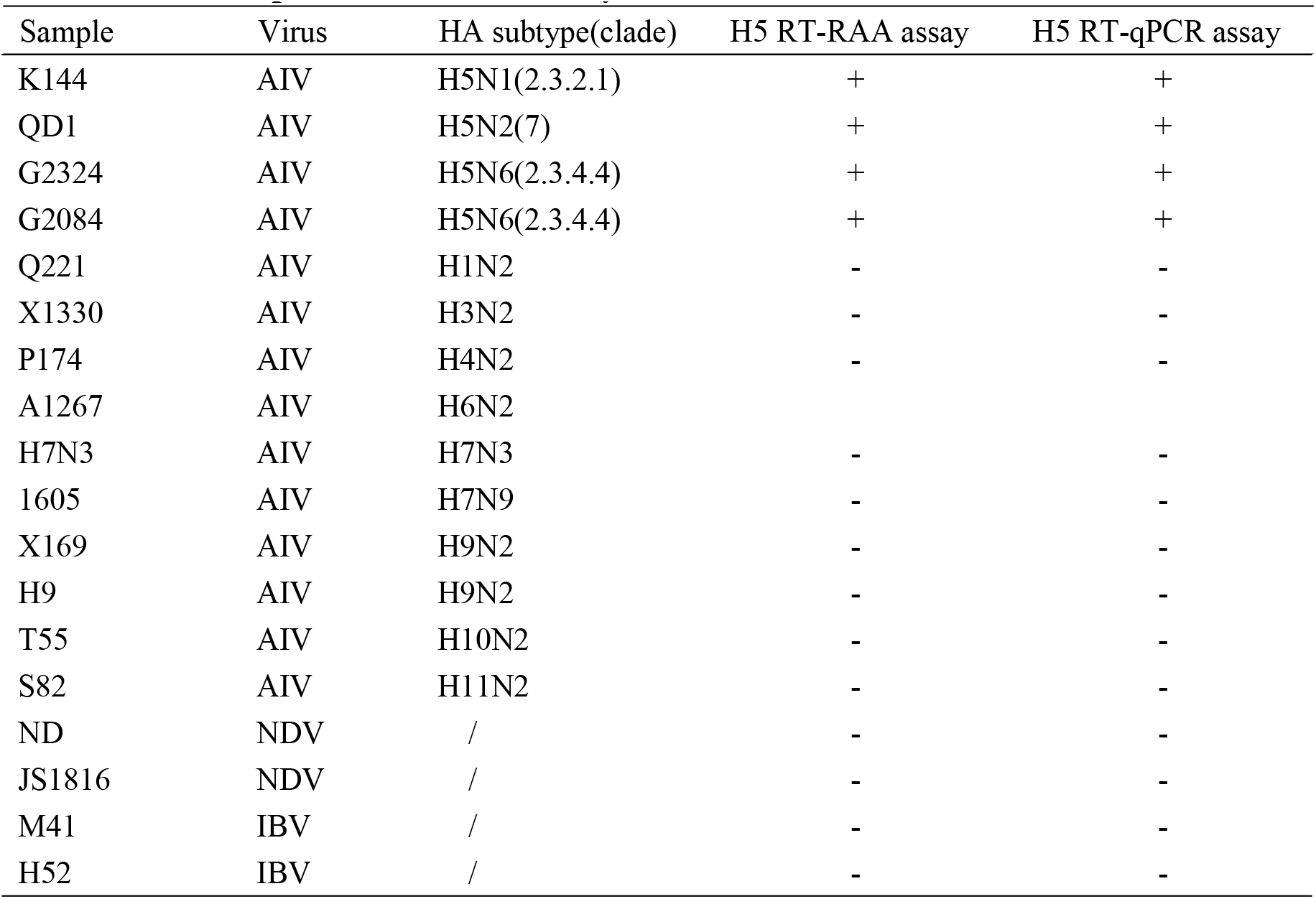
Samples tested in the study. Detection and evaluation of clinical samples by H5 RT-RAA

About 365 Oral-pharyngeal and cloacal swab samples were collected from live poultry market, which were immediately placed into 1mL antibiotic-containing PBS as described above and then stored at −80°C until total nucleic acids were extracted with viral RNA/DNA extraction kit above. The avian clinical samples collected were evaluated the performance of the RT-RAA assay and compared it with the performance of published RT-qPCR assays as described previously for H5 subtype AIV. The primers and probe for the RT-qPCR assays are listed in Table 1[21, 22]. In addition, the positive clinical samples were confirmed by sequence analysis to verify the positive results in conventional PCR.

### Statistical analysis

To determine the RT-RAA detection limit, a probit analysis was performed at a confidence interval of 95%, and the kappa and p values of RT-qPCR and RT-RAA were calculated. In addition, we calculated the sensitivity and specificity of RT-qPCR and RT-RAA for detection in clinical samples of poultry. All statistical analyses were performed in SPSS 21.0 (IBM).

## RESULTS

### Analytical sensitivity of RT-RAA

The sequence of the appropriate H5 RT-RAA primers and exo-probe are listed in Table 1, which has the best amplification efficiency under the same reaction conditions. The detection results of RT-RAA sensitivity assay are shown in Figure 1. The primer and probe combinations designed by the present invention have a RNA concentration of 10^7^ copies /μL, 10^6^ copies/μL, 10^5^ copies/μL, and 10^4^ copies/μL, 10^3^ copies /μL, a fluorescence amplification curve appears. Therefore, the detection limit of H5 RT-RAA assay was 10^3^ RNA copies/μL(Fig 1).

**Fig 1.**
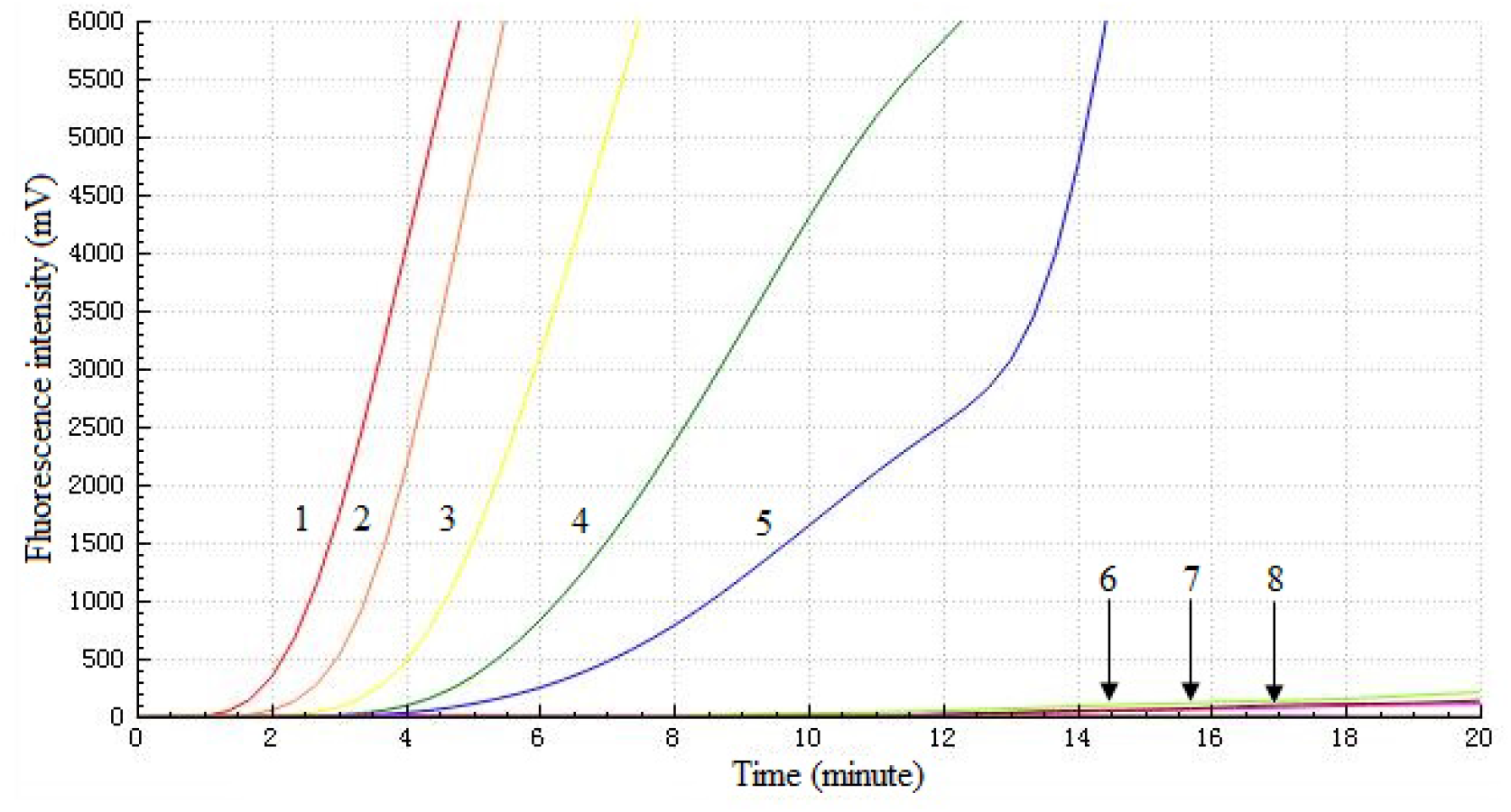
Analytical sensitivity of the H5 RT-RAA assay. A dilution range from 10^7^ to 10^1^ copies per reaction of H5 subtype AIV RNA molecular was, respectively, used to evaluate the detection limit of H5 RT-RAA assay, negative represents negative control. 1—7: 10^7^ copies /μL—10^1^ copies /μL; 8: negative control.

### Analytical specificity of RT-RAA

The results showed that the test group corresponding to the H5 subtype AIV RNA template showed normal fluorescence detection curves, and the other virus test groups and negative control groups did not show amplification curves. Thus, the RT-RAA assay did not cross-react with other subtype AIVs, NDVs and IBDVs and demonstrated high specificity for the detection of H5 subtype AIV(**Fig 2**).

**Fig 2.**
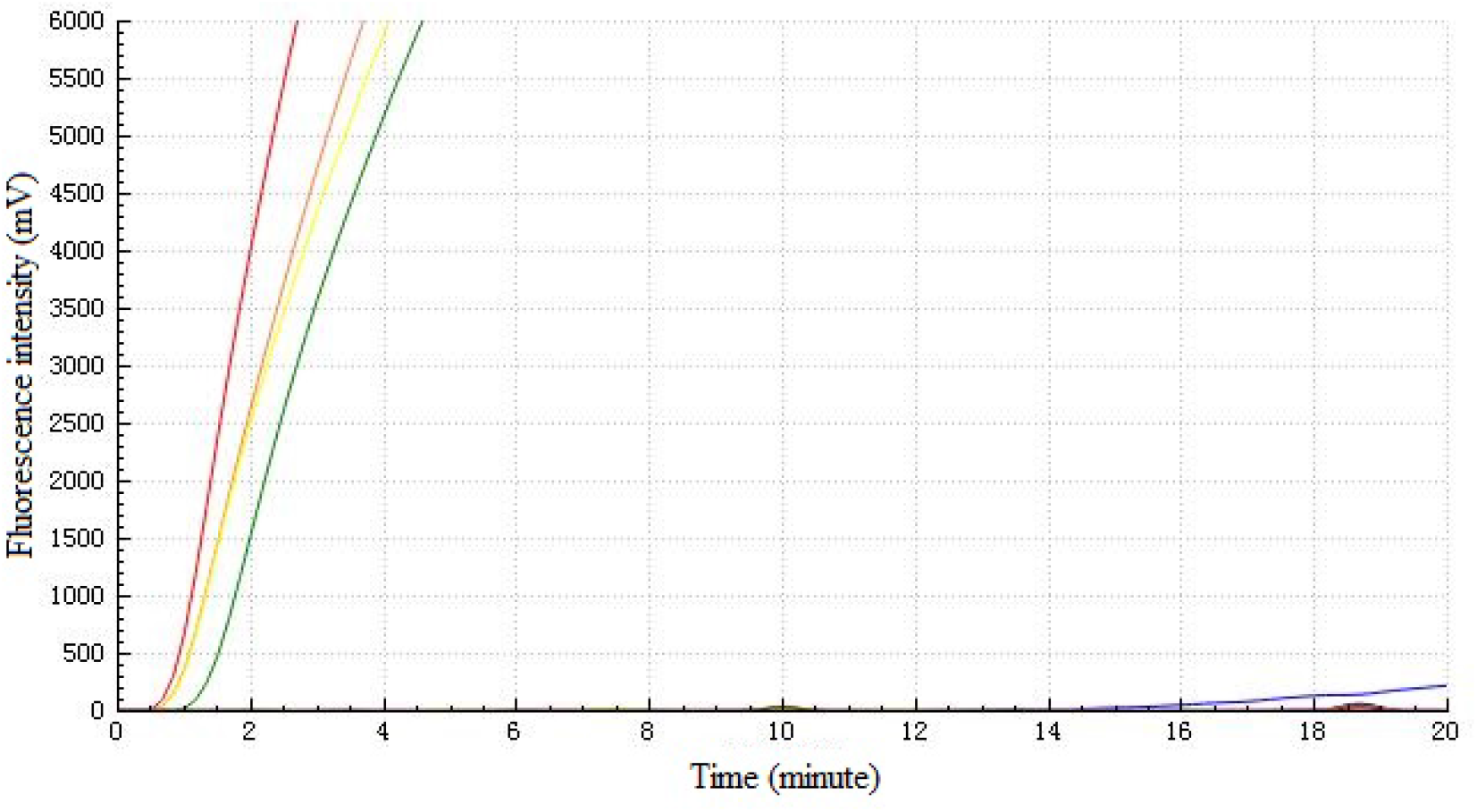
Analytical specificity of the H5 RT-RAA assay. Detection signals were recorded by real-time fluorescence RT-RAA with four samples including H5 subtype AIVs (H5N1, H5N2 and H5N6), while no signals were detected from the fourteen samples including other subtype AIVs, NDVs, IBVs and negative controls.

### Evaluation of the RT-RAA for clinical samples

The 365 conserved avian clinical samples collected from live-poultry markets were tested by RT-RAA assay and compared with RT-qPCR. A threshold cycle (CT) value of 36 was used as the cut-off for a positive result in RT-qPCR. RT-qPCR detected 18 of the 365 samples as positive (4.93%, 18/365), while RT-RAA correctly identified and differentiated 17 positive samples, with a sensitivity of 94.44% (95%CI, 70.63% - 99.71%) and 100% specificity (95%CI, 98.64% - 100%) (Table 3). The κ value for RT-RAA and RT-qPCR was 0.970 (p < 0.001). All the detected positive clinical samples were verified as H5 subtype AIV positive by conventional PCR and sequence analysis.

**TABLE 3.**
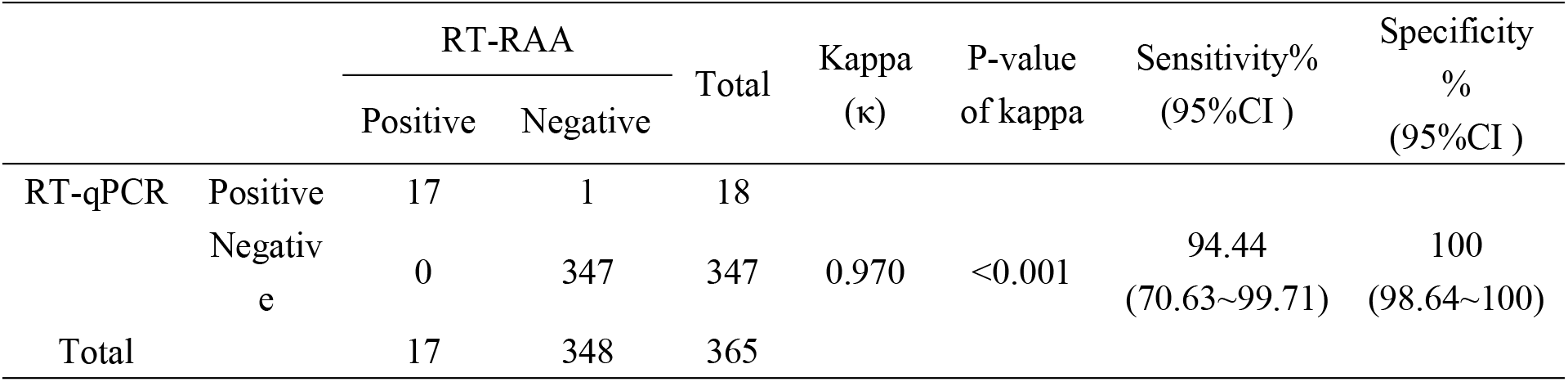
Detection of H5 subtype AIV in avian clinical samples.

## DISCUSSION

The frequent antigen shift and antigen drift of AIV increase the difficulty of AIV detection. Among all the AIV subtypes, highly pathogenic H5 AIV often leads to high morbidity and mortality in poultry. Nowadays, the H5 virus are widely prevalent and of significant concern to the poultry industry and public health in China. Therefore, early detection of AIV and H5 subtype is very necessary in the surveillance and control of AIV outbreaks. Until now, there were many methods used in AIV detection, such as gold immunochromatographic assay [23], microarray [24], immunosensor [25], immune-fluorescence [26] and enzyme linked immunosorbant assay[27]. However, these methods require complex and costly devices and are difficult to perform in the field [28]. So far, only detection methods based on nucleic acid sequence-based amplification [29], RT-LAMP, and RT-RPA [30] have been used for the rapid detection of the AIV in the field. Moreover, as the mutation rate of the H5 subtype AIV accelerates, the previous detection methods may not meet the actual detection needs, some of them fail to subtype AIV while others cannot be applied in early diagnosis owing to inadequate sensitivity [31,33], so a rapid diagnostic method capable of detecting all popular strains is needed.

RAA is a novel isothermal amplification and detection assay requiring only 20 min to complete, compared with approximately 1 and 3 hr, respectively, for RT-qPCR and conventional PCR. In PCR, thermocycling is required for double-stranded DNA separation, primer binding and amplification. There should be no more than 30 amplification cycles for pathogen detection in PCR. RAA could produce a positive signal in as little as 4 min, at half the cost of RT-qPCR [34]. Recombinase-aided amplification can be carried out using a portable device with no complicated processes, while the instruments for RT-qPCR and conventional PCR are much more expensive. Recombinase-aided amplification can also use reverse transcriptase and a fluorescent probe system to detect RNA amplicons in real time [35]. Until now, RAA has only been used for detecting human pathogens, including Salmonella, respiratory syncytial virus [36], coxsackievirus [34], hepatitis B virus [37] and Schistosoma japonicum-specific gene fragments[38] and there have been no previous reports of the use of RAA for detecting H5 subtype AIV[39].

In this study, an H5 RT-RAA assay was created, which indicating the potential value of our method in application of early detection and rapid diagnosis of the H5 subtype AIV. In the detection of experimental and clinical samples, this method showed higher sensitivity along with high efficiency. The results showed that the RT-qPCR detected 18 of the 365 samples as positive (4.93%, 18/365), while RT-RAA correctly identified and differentiated 17 positive samples, with a sensitivity of 94.44% (95%CI, 70.63% - 99.71%) and 100% specificity (95%CI, 98.64% - 100%) (Table 3). The κ value for RT-RAA and RT-qPCR was 0.970 (p < 0.001). Moreover, the clinical swab samples detected as positive in RT-RAA were also certified as positive by PCR detection and sequencing. According to the primer/exo-probe design method and the sample detection results, the RT-RAA method was considered to be suitable for detecting almost all H5 subtypes. To the best of our knowledge, this is the first RT-RAA method for the detection of multiple H5 clades in AIV.

In conclusion, the RT-RAA method established in our study can quickly and accurately identify H5 subtype AIV, including the current epidemic strains, which meets the need for H5 subtype AIV testing, It is expected that this RT-RAA for detection of emerging H5 subtype AIV rapidly will be applied in surveillance of clinical samples in field experiments and provide a powerful and valuable tool for the control of H5 subtype AIV.

## Acknowledgement

This work was supported by the National Key Research and Development Program of China (2017YFC120050)

## Author Contributions

**Conceptualization**: Suchun Wang, Yang Li, Kai-Cheng Wang, Hualei Liu and Baoxu Huang.

**Data Curation**: Suchun Wang, Kai-Cheng Wang.

**Formal Analysis**: Suchun Wang, Yang Li.

**Funding Acquisition**: KaiCheng Wang, Baoxu Huang.

**Investigation**: Suchun Wang, Yang Li, Nan Jiang, Fuyou Zhang, Qingye Zhuang, Guangyu Hou.

**Methodology**: Suchun Wang, Yang Li, KaiCheng Wang.

**Project Administration**: KaiCheng Wang.

**Resources**: Guangyu Hou, Qingye Zhuang, Lijian Jiang, Jianmin Yu, Chenglong Zhao, Xiaohui Yu and Liping Yuan.

**Supervision**: KaiCheng Wang.

**validation**: Suchun Wang, Yang Li, Nan Jiang, Fuyou Zhang.

**Visualization**: Suchun Wang, Yang Li.

**Writing – Original Draft Preparation**: Suchun Wang, Yang Li.

**Writing – Review & Editing**: Suchun Wang, Yang Li, Kai-Cheng Wang, Hualei Liu, Baoxu Huang.

